# Assessing extracellular vesicles from bovine mammary gland epithelial cells cultured in FBS-free medium

**DOI:** 10.1101/2021.09.29.462434

**Authors:** G. Silvestrelli, S.E. Ulbrich, M.D. Saenz-de-Juano

## Abstract

**Aim:** Mammary gland extracellular vesicles (EVs) are found in both human and livestock milk. Our knowledge of the role of EVs in the mammary gland development, breast cancer and mastitis derives mainly from *in vitro* cell culture models. However, a commonly shared limitation is the use of foetal bovine serum (FBS) as a supplement, which naturally contains EVs. For this reason, the purpose of the study is to establish a novel tool to investigate mammary gland EVs *in vitro* and in an FBS-free system.

**Methods:** Primary bovine mammary epithelial cells (pbMECs) and a mammary gland alveolar epithelial cell line (MAC-T) were cultured in a chemically defined EV-free medium. To find a reliable EVs isolation protocol from a starting cell conditioned medium (10 mL), we compared eight different methodologies by combining ultracentrifugation (UC), chemical precipitation (CP), size exclusion chromatography (SEC), and ultrafiltration (UF).

**Results:** The medium formula sustained both pbMECs and MAC-T cell growth and did not alter MAC-T cell identity. Transmission electron microscopy revealed that we obtained EV-like particles in five out of eight protocols. The cleanest samples with the highest particles amount and detectable amounts of RNA were obtained by using UF-SEC-UC and UC-SEC-UC.

**Conclusion:** Our chemically defined, EV-free medium sustains the growth of both pbMECs and MAC-T and allows the isolation of EVs that are free from any contamination by UF-SEC-UC and UC-SEC-UC. In conclusion, we propose a new culture system and EVs isolation protocols for further research on mammary epithelial EVs.

## INTRODUCTION

Mammary gland extracellular vesicles (EVs) participate in many physiological processes of the mammary gland such as development [1, 2] and regulation of epithelial cells polarity [3]. They are likewise involved in pathological conditions, including mastitis [4, 5] and breast cancer in primis, where EVs are implicated in its onset [6], metastasis [7] and drug resistance [8, 9]. Mammary gland EVs are also found in the milk of human [10] and many livestock species such as cattle [11, 12], buffalo [13] and goat [14].

Our knowledge of the molecular mechanisms underlying EVs biogenesis, release, uptake, and effect on recipient cells, derives mainly from in vitro cell culture models. Many studies on human cell lines have investigated the role of EVs in mammary gland communication in the context of cancer [9, 15, 16] or the maintenance of epithelial cells polarity [3]. In mice, EVs play an important role in mammary gland development [3] and involution [2], and EVs promote the recognition and clearance of apoptotic bodies from macrophages.

While most of the published studies focus on humans and mice, only a few studies investigated EVs directly in bovine mammary gland culture models, from primary cells [17] or cell lines [18]. Bovine milk EVs are heterogenous and comprehend many EVs subtypes including exosomes (40-100 nm) and microvesicles (100-1000 nm). In addition, milk EVs are biologically active, they are transferred to the newborn and are taken up from intestinal cells [11]. To date, their role is still not known in cell-cell communication within the alveolus of the mammary gland. Besides, the origin of milk EVs is not fully clear, despite recent publications suggesting that they are produced by the milk-secreting epithelial cells, the lactocytes [19, 20]. Of note, autocrine and paracrine communication within the bovine mammary gland alveolus is important in all its developmental steps [21], and, in this frame, EVs might also participate in this intense communication.

Cell culture systems allow more options and flexibility to investigate EVs mediated communication as compared to in vivo systems. However, they usually share a limitation, namely the use of foetal bovine serum (FBS) as a supplement. FBS naturally contains EVs that can interfere with the experimental setting and outcome [22], therefore many studies use commercial or home-made EV-depleted FBS, also in bovine MECs-EVs research [23]. However the removal of EVs from FBS is never complete [22] and alters the FBS effect on cells, affecting cell growth [24], differentiation [25], and response to pathogens [26, 27]. A period of 24-48 hours of starvation from FBS affects the culture conditions as well, leading to cell cycle arrest [28] and reducing the background production of cytokines [29], which can cause misinterpretation of cells phenotype and experimental outcome. For these reasons, it is recommended to culture cells directly in FBS-free systems and chemically defined media. Besides the type of medium, it is critical to collect enough EVs for downstream analyses. For this reason, the volume of the starting material usually ranges between 20 and 250 mL [30-32]. Another factor affecting the yield and purity of the EVs sample is the isolation procedure [33].

Few cell lines from the bovine mammary gland exist, with BME-UV1 [34] and MAC-T [35] as the most commonly known to date. The use of pure epithelial cell lines prevents contamination from fibroblasts often occurring in primary cultures, which can be also avoided by pre-plating [36], short trypsinization and isolating epithelial cells from milk [37]. MAC-T cells are considered a proper model for mammary gland development and lactation [38], as they maintain the capacity to express milk proteins [39] and are responsive to hormonal stimulation [34]. However, cell lines tend to lose the phenotypes of the original tissue [40] and they are set to grow in specific culture conditions, therefore the change of culture conditions can also alter their characteristics.

To gain novel insights on the EVs local communication of the bovine mammary gland in vitro models compiling the guidelines for EVs research are in need. For this reason, in the current study we i) developed a chemically defined FBS-free culture that supports the growth of both primary MEC and MAC-T cell line and ii) compared eight different EVs isolation protocols from only 10 mL of conditioned medium regarding quantity and size distribution of EVs for further downstream analyses.

## METHODS

### Primary cells isolation and culture

Mammary glands from lactating cows were collected at a local slaughterhouse post mortem and transported on crushed ice to the lab. Only mammary glands that did not display fibrosis, abnormal cell growth, and signs of mastitis (e.g. redness or hardness) were used to set the cultures. Tissue pieces of ∼10 g were pooled from 2 to 4 cows each and washed in ethanol 70%, then in cold PBS (Gibco, Thermo Fisher, USA) with antibiotics. Tissues were further minced in ∼2 mm3 pieces and washed 6 times in cold PBS with antibiotics. They were digested in Collagenase IV (Sigma-Aldrich, USA) 0,5 mg/mL, Dispase II (Sigma) 0,5 mg/mL, Insulin 5 µg/mL (Sigma), antibiotics and antimycotics in HBSS (Gibco) buffer for 2 hours at 37°C while gently shaking. The suspension was filtered through a metal mesh to remove larger tissue pieces and then centrifuged at 500 xg for 5 minutes. The pellet was washed twice in PBS and cells were seeded on Nunclon Delta surface dishes (Thermo Fisher) and kept at 37.5 °C 7% CO2. For the pre-plating, freshly isolated cells were plated and let for 1 hour in the incubator, then the medium and cells that were not attached yet were transferred into a new plate. The medium was changed every 2-3 days until 80% confluence, cells were then subpassaged every 3-4 days. Cells were kept in DMEM/F12 (Gibco), containing Gentamicin 50 µg/mL (Sigma), Amphotericin B 2,5 µg/mL (Sigma) and supplemented with a) FBS 10%, or b) B27 1:50 (Gibco), 5 µg/mL Insulin (Sigma), 5 µg/mL Hydrocortisone (Sigma), Estradiol (E2) 100 nM (Sigma), Progesterone (P4) 300 pM (Sigma) and 5 ng/mL Epidermal Growth Factor (EGF) (Sigma). The latter medium is referred to as FBS-free medium. When plates were coated, rat tail collagen I (Invitrogen, USA) at the final concentration of 6 or 10 µg/cm2, or laminin (Sigma) 1 or 2 µg/cm2 was used. For the growth curve, 6 × 105 cells were seeded on 6-well multiwell plates and counted every two days from day 4 to day 14. To evaluate the growth, 6 × 105 cells were seeded on 6-well multiwell plates and counted every two days from day 4 to day 14 using Trypan blue (Sigma) and a Neubauer chamber.

### MAC-T cell line

MAC-T cells were kindly provided by Olga Wellnitz from the Vetsuisse Faculty of the University of Bern (Switzerland). Cells were cultured in FBS 10% or FBS-free medium and were passaged every 3-4 days. After 2 weeks of adaptation in the FBS-free medium, they were lysed in TRIzol (Thermo Fisher) for checking cell type markers (keratin 18, keratin 14, vimentin) expression. For growth rate evaluation, 1 × 105 cells were plated and counted at 80% confluence for two consecutive passages (referred to P1 and P2). The growth rate was calculated as ln(cells t0/cells t1)/t1-t0.

### RNA isolation and retrotranscription

Once cells reached 80-90% confluency, they were washed in PBS and lysed in TRIzol (Thermo Fisher), followed by phenol-chloroform RNA isolation. DNA was removed by DNAse I treatment (Sigma). For each sample, 500 ng of total RNA were reverse transcribed using the GoScript Reverse transcription system kit (Promega, USA), following manufacturer instructions and cDNA samples were then stored at - 20°C until further use.

### Gene expression analysis

Table 1 shows the primer pairs, actin, GAPDH, and histone H3 were used as reference genes. For the RT-qPCR the Kappa Mix (Sigma) was used and the primers concentration was 10 µM. The amplification was performed using 500 ng of cDNA and the following amplification program: 95°C 3’ (x 1), 95°C 4’’, 60°C 20’’, 95°C 10’ (x40). Relative gene expression analysis was performed using the 2-ΔΔCt method [40].

**Table 1.**
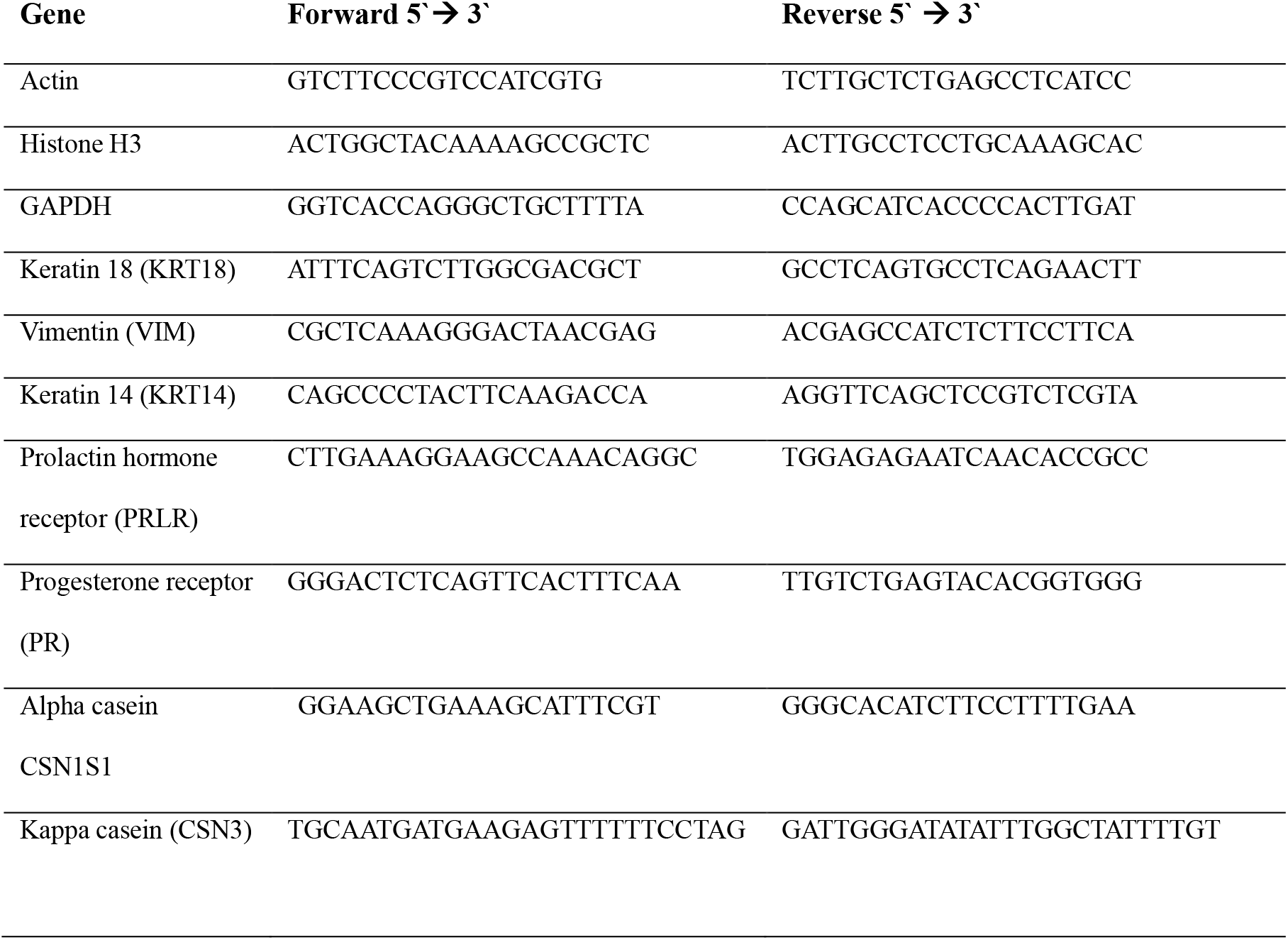
List of the designed forward and reverse primer pairs, KRT14 and PRLR are from Finot et al. 2018 [41]

### Differential centrifugation (DC)

The conditioned medium (CM) from cells cultured in FBS-free conditions was collected at 80-90% confluency. The CM was differentially centrifuged at 300 xg, 1000 xg, 12000 xg 10 min at 4°C to remove dead cells, cells debris and apoptotic bodies, respectively. Immediately, the CM was frozen at -80°C until the EV isolation. We combined different methods to isolate EV from 10 mL of conditioned medium, testing in total 8 routes which are schematized in figure 1. We started from 0,5 mL for SEC-UC.

**Figure 1.**
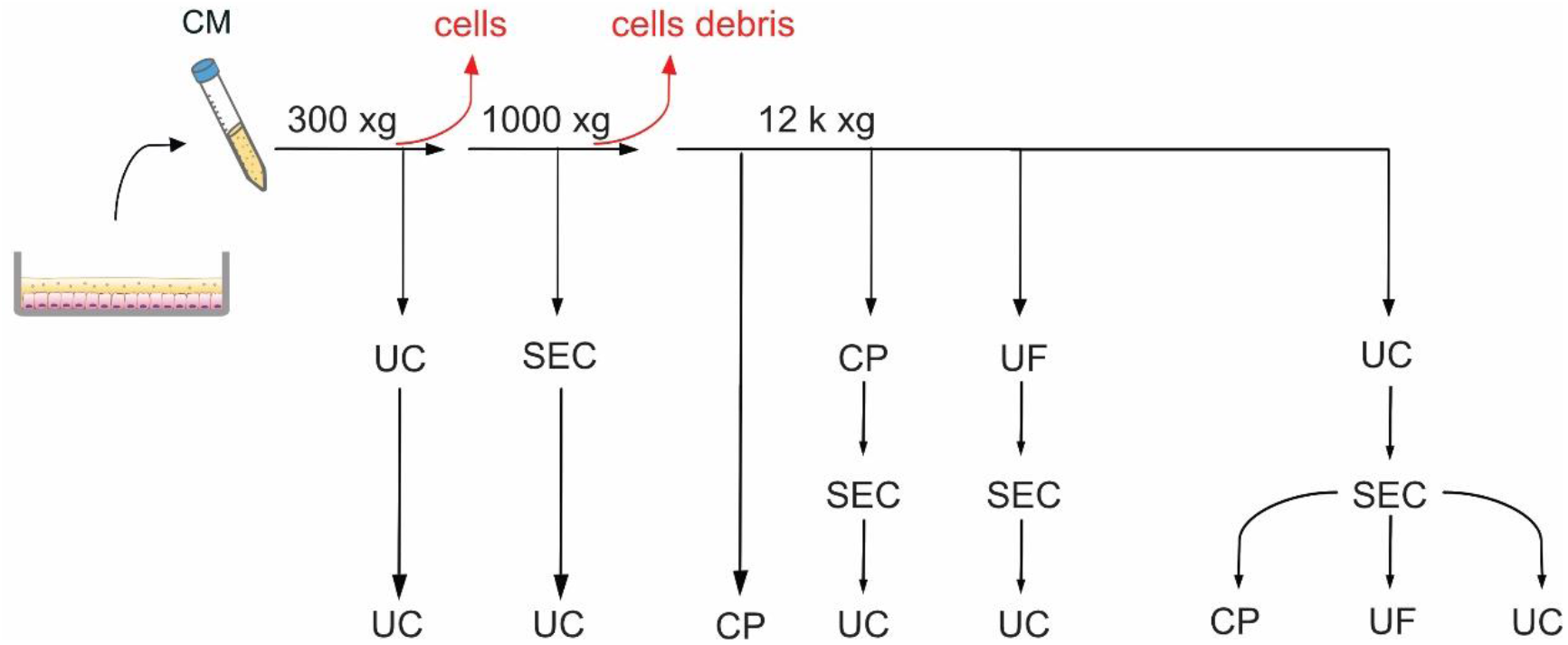
Schematic representation of the 8 protocols (routes) tested. UC: ultracentrifugation, UF: ultrafiltration with Amicon tubes, CP: chemical precipitation with miRCURY (Qiagen), SEC: size exclusion chromatography with qEV columns (IZON).

### Ultracentrifugation (UC)

The CM was thawed on ice and transferred into 13.5 mL Ultra-Clear ultracentrifuge tubes (Beckman Coulter, USA). Samples were spun at 120k xg for 2 hours or 200k xg for 70 min at 4°C in a Beckman Coulter Ultracentrifuge Optima XE-90, using a Type 50.2 Ti rotor. The pellets were resuspended in 0.22 µm filtered PBS for further ultracentrifugation, or size exclusion chromatography, and stored at -80°C.

### Ultrafiltration (UF)

To perform UF, 10 mL of CM or qEV fractions 6-10 diluted in PBS were loaded on 15 mL Amicon Tubes with a cut-off of 100 kDa (Merck, Germany) and spun for 1 hour RT at 5000 xg on a FA-45-6-30 fixed angle rotor. The concentrate was transferred into a new tube and either frozen or loaded onto a qEV column (IZON Science Ltd, New Zealand).

### Precipitation with miRCURY (Qiagen)

To chemically precipitate the EVs, we used the mirCURY kit (Qiagen, Germany) for cell culture medium, following the manufacturer instructions. Briefly, the CM after differential centrifugation or from size exclusion chromatography was spun at 3000 xg at 4°C for 5 minutes to remove debris and cryo-precipitated particles. The supernatant was transferred into a new tube where 0.4 of sample volume of buffer B was added. After vortexing, the tubes were left on ice for 1 hour and then spun at 20°C at 3200 xg for 30 minutes. The precipitate was resuspended in 100 µL of resuspension buffer and either snap-frozen or loaded onto qEV columns.

### Size exclusion chromatography (SEC)

We performed size exclusion chromatography with qEV 70 nm classic columns (IZON). The columns were first equilibrated at room temperature (RT) with 10 mL of 0.22 µm filtered PBS (Gibco), then 500 µL of CM or EV resuspension from UC or UF was loaded on the column and eluted in filtered PBS. The flow-through was collected in 500 µL fractions and in each fraction the protein concentration was measured by the A280 at the Nanodrop. Fractions 6-10 were pulled together for further concentration.

### Tunable resistive pulse sensing (TRPS) measurements

All measurements were conducted using a qNano Gold (IZON) and NP150 polyurethane nanopores (IZON), that detects particles with a diameter ranging from 70 to 420 nm. Filtered PBS (Gibco) was used as an electrolyte buffer and CPC100 (IZON) as calibration particles. Analyses were performed with Izon Control Suite v.3.3.

### Transmission Electron Microscopy (TEM)

The EVs visualization was performed at the Scientific Center for Optical and Electron Microscopy (ScopEM) service of ETH Zurich. Briefly, 3 µL of the vortexed dispersion were placed on glow discharged carbon-coated grids (Quantifoil, D) for 1 minute. Negative contrast staining was done in 2% sodium phosphotungstate pH 7.2 for 1 second, followed by a second step for 15 seconds. Excess moisture was drained with filter paper and the imaging of the air-dried grids was done in a TEM Morgagni 268 (Thermo Fisher) operated at 100 kV. For each experimental group, two replicates were analysed.

### Proteins isolation and Western blot analysis

The pellet of freshly isolated EVs from 45 mL of medium was immediately lysed in RIPA buffer plus protease inhibitors and then stored at -80°C. To each sample, 4x Laemmli buffer (Biorad, USA) was added and heated for 10 min at 95°C. Between 1-3 µg of proteins (for bMEC EV, milk EVs, and cell lysates) were run on 12% polyacrylamide gel and then total protein was evaluated at the Chemidoc running the stain-free program. Proteins were transferred using a TransTurbo transfer pack (Biorad) with a TurboBlot (Biorad). Membranes were blocked 1 hour in skim milk 5% TBS-Tween buffer (TBST, Bio-Rad, 0.05% Tween 20), and incubated overnight with the primary antibodies diluted in blocking buffer: anti-TSG101 (1:250, PA531260, Thermo scientific) 1:250, anti-calnexin (1:2000, ab75801, Abcam, UK) anti CD9 (1:250, MM2/57, Biorad). After 3 washes in TBST, membranes were incubated for 1 hour at RT with secondary antibodies (Santa Cruz Biotechnology, USA) and StrepTactin-AP Conjugate (Biorad) at the concentration of 1:10000, eventually incubated with Clarity Western ECL Substrate (Biorad) for chemiluminescent signal development.

### RNA isolation

Total RNA including microRNA (miRNA) were isolated with the miRNeasy MicroKit (Qiagen, Germany). The concentration was measured with Quantus™ Fluorometer and the QuantiFluor® RNA System kit (Promega). The length from RNA fragments was evaluated using the Agilent Pico Kit and the Agilent 2100 BioAnalyzer (Agilent Technologies).

### RNA retrotranscription and RT-qPCR

For each sample, 10 ng of RNA were retrotranscribed and pre-amplified using the TaqMan™ Advanced miRNA assays (Life Technologies), following manufacturer instructions. Then, we performed an RT-qPCR using the kit TaqMan™ Advanced miRNA assays (478575_mir assay), targeting the miR-let7a-5p. The amplification program was: 95°C 30’’ (x 1), 95°C 5’’, 60°C 30’’ (x40).

### Statistical analysis

To evaluate the gene expression of mammary epithelial markers we performed the non-parametric Friedman test and Dunn’s multiple comparisons tests using GraphPad Prism version 8.2. The differences were considered significant when p <0.05. Unless stated, all values are given as mean ± standard deviation (SD).

## RESULTS

### Evaluation of cells growth in FBS-free medium

From the isolation (day 0) to 80% confluency, primary bMECs took in average 18.5 ± 4.07 days when grown in FBS-free medium. Cells grew from ∼1×10^5^ at day 4 to ∼1.1×10^6^ (day 12) (figure 2a). Coating the plates with collagen I or laminin neither improved cell attachment nor growth during the first days (supplementary figure 1a), therefore, we kept plating on uncoated wells. The pbMECs were cultured over three passages in FBS-free medium. MAC-T cells grew in FBS-free medium as well, and the growth rate did not change compared to FBS containing medium (p>0.05, table 2).

**Table 2.** Growth rates of MAC-T grown in FBS containing or FBS-free medium in two consecutive passages. Data are expressed as mean ± SD of 3 independent experiments. Mann-Whitney test was performed

| MAC-T cells | Passage 1 [h<sup>-1</sup>] | Passage 2 [h<sup>-1</sup>] |
| --- | --- | --- |
| FBS | 0.011 ± 0.006 | 0.023 ± 0.003 |
| FBS-free | 0.022 ± 0.015 | 0.012 ± 0.005 |

**Figure 2.**
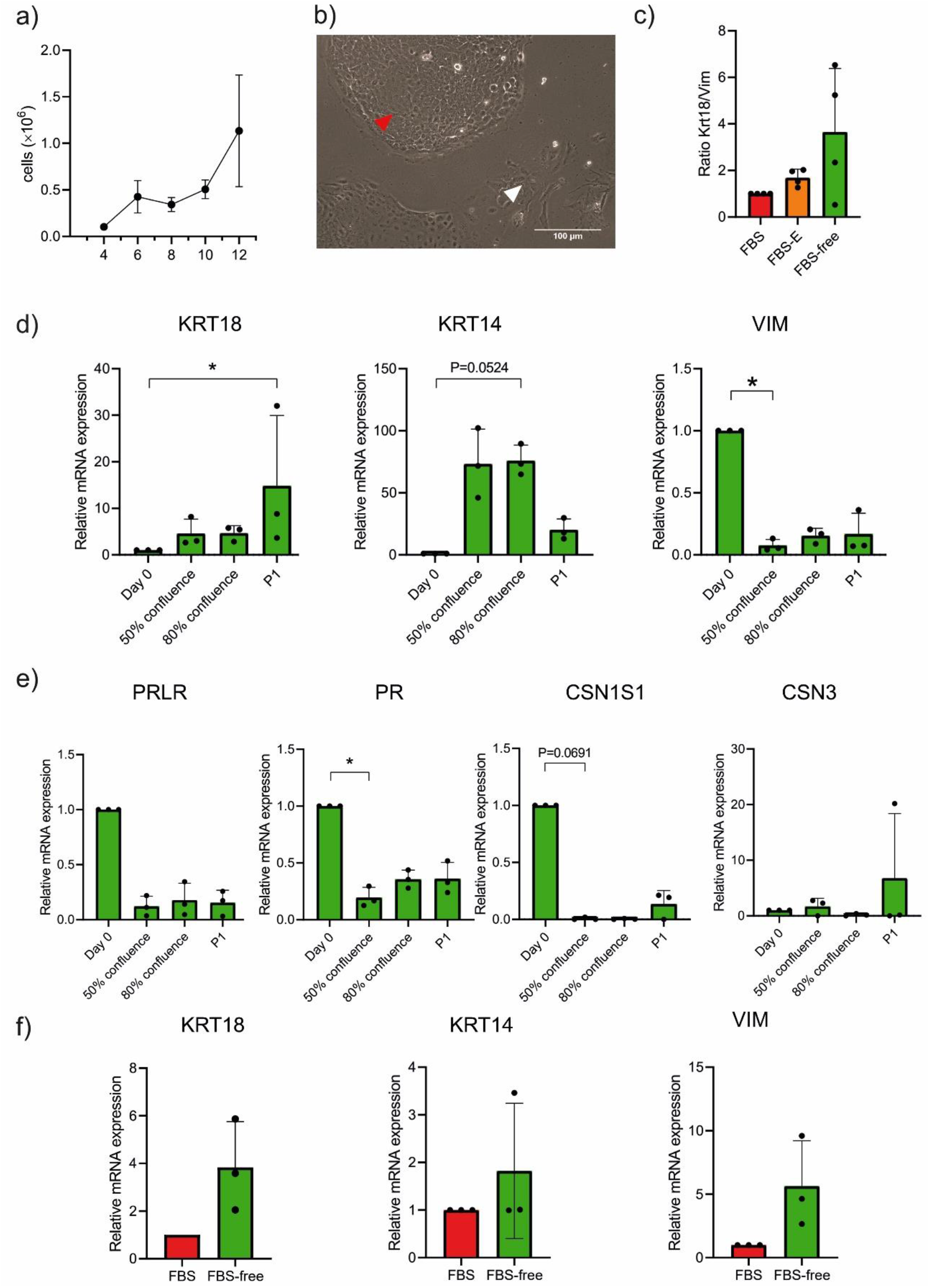
The pbMEC and MAC-T properties in FBS-free medium. a) Cell counts from day 4 to day 12 in 6-wells multiwell culture dishes; b) Epithelial cells population (red arrow), and fibroblast-like cells (white arrow); c,d) Cell-type (c) and differentiation (d) markers at the isolation day (day 0), at 50% and 80% confluency, and at 80% confluency at the first sub-passage (P1): keratin 18 (KRT18), keratin 14 (KRT14) for epithelial identity, vimentin (VIM) as fibroblastic marker, prolactin hormone receptor (PRLR), progesterone receptor (PR), casein alpha (CSN1S1) and casein kappa (CSN3) coding genes as differentiation markers; f) Keratin 14 and 18 and vimentin mRNA expression for MAC-T cells cultured in FBS 10% or FBS-free medium. The mRNA expression data are depicted as 2-ΔΔCt [42], values are mean ±SD of 4 independent replicates, Kruskal-Wallis and Dunn’s multiple comparisons tests was performed. The differences were considered significant when p <0.05.

### Evaluation of mammary gland gene markers expression in FBS-free medium

The primary cultures were initially not pure and were composed of a mixed population of fibroblasts (figure 2b, white arrow) and epithelial cells (figure 2b, red arrow). We evaluated the epithelial cells enrichment as indicated by the gene expression ratio between keratin 18 (an epithelial marker) and vimentin (a fibroblastic marker) in pbMECs cultured in i) FBS 10% ii) FBS 10% + pre-plating iii) FBS-free medium. The enrichment of epithelial cells increased 1.7-fold by pre-plating and 3.4-fold in FBS-free medium (figure 2c), compared to pbMECs in FBS without pre-plating. Even if not statistically significant (p=0.1377, FBS-free medium vs FBS 10%), we observed that some cultures reacted to the FBS-free medium by enriching the population in epithelial cells, without the need of pre-plating. Coating the plate with neither collagen I nor laminin significantly affected the enrichment (p>0.05, supplementary figure 1b).

Subsequently, we evaluated the gene expression of cell type and differentiation markers at day 0 (isolation day), 50% and 80% confluence after day 0, and at 80% confluence after the first sub-passage (P1). The mRNA abundance of the epithelial markers keratin 14 (KRT14) and 18 (KRT18) increased at 80% confluency after the isolation day and at P1, respectively (p<0.05, figure 2d). On the other hand, the expression of the fibroblastic marker vimentin (VIM) decreased already at 50% confluence after day 0 (p<0.05, figure 2d). The expression of the functional marker progesterone receptor (PR) significantly decreased already at 50% confluence (p<0.05, figure 2e), while casein alpha (CSN1S1) and prolactin hormone receptor (PRLR) tended to decrease over time (p>0.05, figure 2e). Casein kappa (CSN3) did neither significantly change, but the trend was rather to increase in some replicates (figure 2e).

MAC-T cells cultured in FBS-free medium presented as a homogeneous population and did not morphologically differ from the FBS-containing environment (supplementary figure 1c). Gene expression of KRT14, KRT18 and VIM neither changed (p>0.05, figure 2f), indicating that the FBS-free medium did not alter the cell-type composition.

### Evaluation of EVs isolation methods from pbMECs and MAC-T conditioning medium

As figure 3a shows, the conditioning FBS-free medium alone did not contain any vesicles, ruling out any contamination of EVs from the medium used. According to the TEM, we obtained particles from all routes except from SEC-UC (figure 3c).

**Figure 3.**
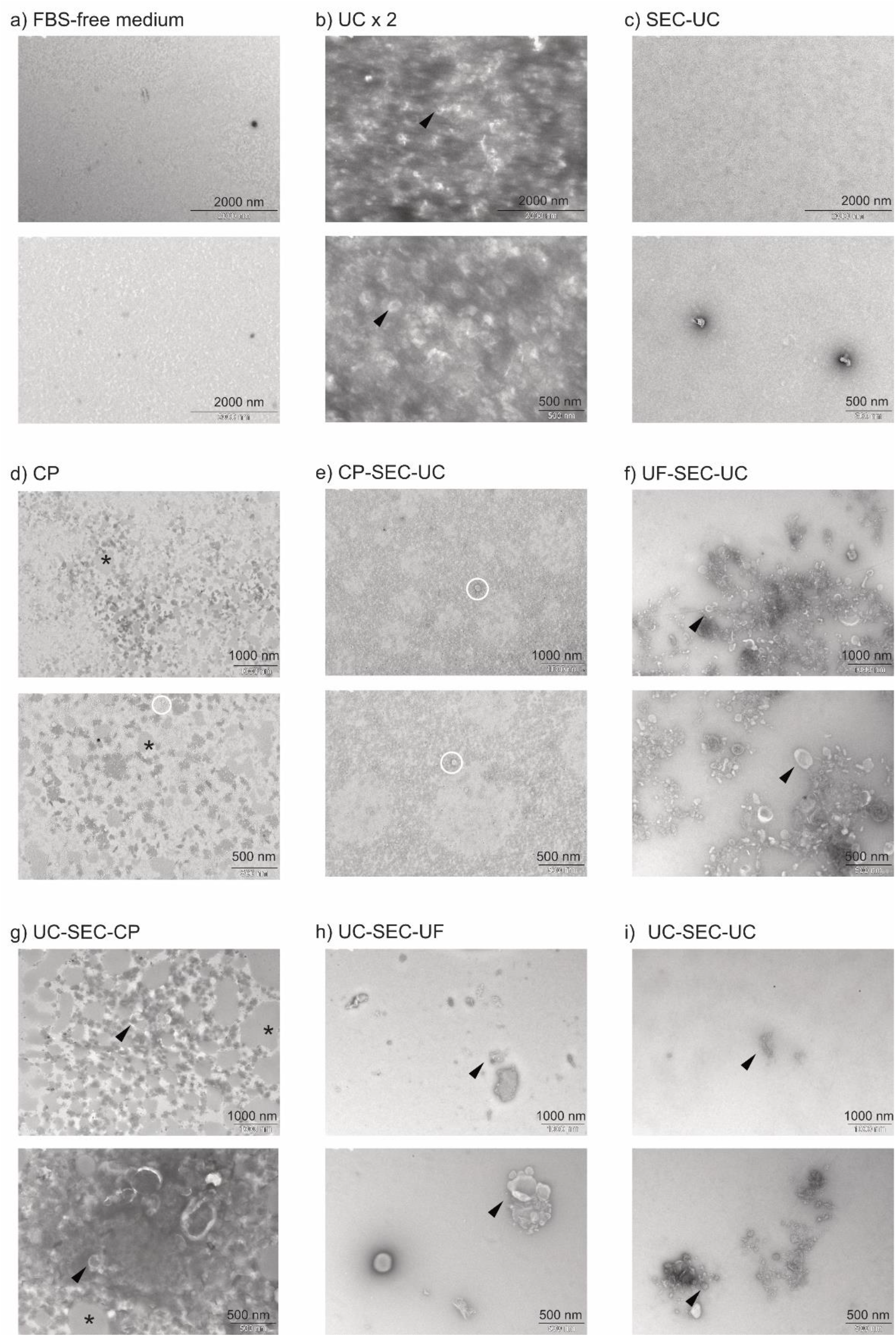
TEM pictures from FBS-free medium (a) and EVs isolated with UC x2 (b), SEC-UC (c), CP (d), CP-SEC-UC (e), UF-SEC-UC (f), UC-SEC-CP (g), UC-SEC-UF (h), UC-SEC-UC (i). Black arrow: cup-shaped particles (vesicles); white circles: non-vesicles particles; black asterisks: aggregates

Single or aggregates of cup-shaped particles exhibiting the typical EV morphology at TEM [41, 42] were observed following UCx2, UF-SEC-UC, UC-SEC-UC, UC-SEC-UF, UC-SEC-UC, (figure 3b,f,g,h and i; black arrows). The observed structures were heterogeneous in size, ranging from 50 to 500 nm. We also observed light grey aggregates in CP, CP-SEC-UC and UC-SEC-CP (figure 3d, e and g; white asterisk), likely due to lipid aggregates [43, 44], while round non-vesicles [45] were observed from CP, CP-SEC-UC and UF-SEC-UC (figure 3d, e and f; black asterisk).

Table 3 summarizes particle concentration and size measured by tunable resistive pulse sensing (TRPS). The particle concentration ranged from 108 to 1011 particles/mL and a particle size between 67 and 781 nm. We did not detect any particles if following SEC-UC and UC-SEC-UF.

**Table 3.** Summary of TRPS measurements. The data come from one replicate.

| Route | Pellet<br>resuspension<br>volume [μL] | Raw<br>concentration<br>[particles/mL] | Size range | Mean particle size<br>[nm] ±SD | Particle rate<br>[particles/min] |
| --- | --- | --- | --- | --- | --- |
| UCx2 | 30 | 4.39E+11 | 67-512 | 119 ± 39.7 | 192.8 |
| SEC-UC | 30 | - | - | - | - |
| CP | 100 | 2.95E+09 | 108-781 | 195 ± 70.1 | 119.5 |
| CP-SEC-UC | 100 | 4.99E+08 | 123-708 | 210 ± 89.4 | 20.02 |
| UF-SEC-UC | 30 | 5.30E+10 | 70-473 | 139 ± 36.5 | 277.4 |
| UC-SEC-CP | 30 | 1.47E+10 | 81-564 | 151 ± 70.0 | 701.2 |
| UC-SEC-UF | 150 | - | - | - | - |
| UC-SEC-UC | 30 | 6.87E+09 | 77-559 | 158 ± 60.8 | 327.0 |

The EVs collection routes UCx2, UF-SEC-UC, UC-SEC-CP and UC-SEC-UC yielded a higher particle concentration, in line with the TEM results. The TRPS however did not discriminate between the different kinds of particles [46], such as between vesicles and the undefined aggregates in UC-SEC-CP. Therefore, since UF-SEC-UC and UC-SEC-UC had the cleanest background free of undefined aggregates (figure 3f, i), we selected these two routes to continue with. We obtained similar size distributions (figure 4a) across replicates as well as concentration (p>0.05, figure 4b). The last SEC fractions after UF had lower protein concentration than after UC, showing that UF as a first step cleaned better the samples from contaminating proteins (figure 4c).

**Figure 4.**
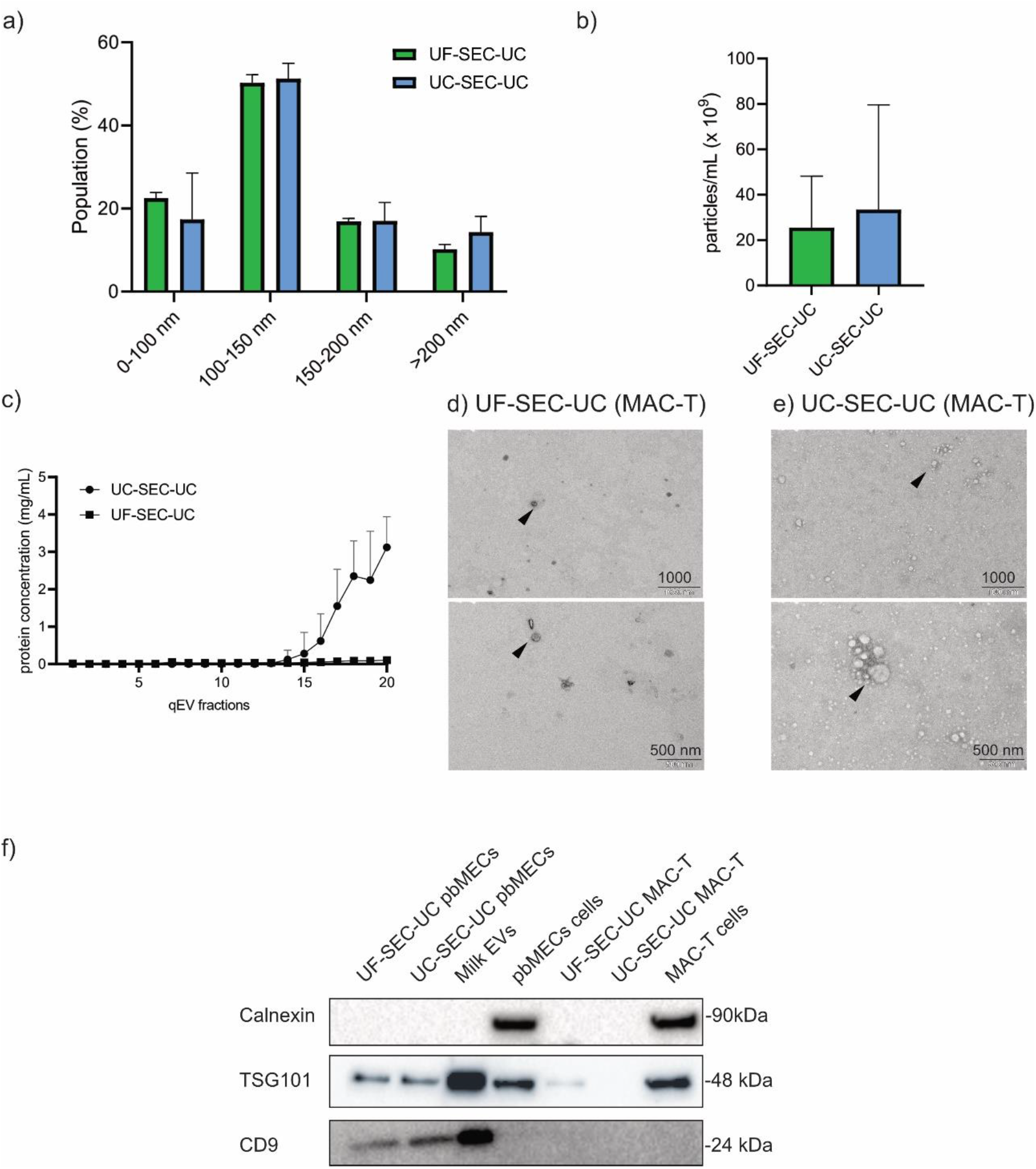
a) Size ranges and b) particle concentration of EVs from UF-SEC-UC and UC-SEC-UC in pbMEC conditioning medium; c) protein concentration of the single fractions from SEC from pbMECs conditioning medium; d-e) TEM pictures after UF-SEC-UC and UC-SEC-UC from MAC-T conditioning medium c) Western blot of pbMECs and MAC-T EVs pellet, whole cells lysates were used as a positive control for calnexin, EVs from milk as a positive control for EV markers. Full-length blots are presented in Supplementary figure 2. In a-c) values are mean ±SD of three replicates.

When isolating EVs from MAC-T with UF-SEC-UC and UC-SEC-UC, we obtained in both routes cup shaped particles as shown with TEM (Figure 4d). The TRPS size and concentration of the isolated particles were 1.63E+E09 and 2.21E+10 particles/mL from UF-SEC-UC and UC-SEC-UC respectively, and similar size ranges (table 4).

**Table 4.** Summary of particle concentrations and diameters measured by TRPS for the routes UF-SEC-UC and UC-SEC-UC in MAC-T.

| Route | Concentration<br>[particles/mL] | Diameter range [nm] | Average particle size [nm]<br>±SD |
| --- | --- | --- | --- |
| UF-SEC-UC | 2.24E+10 | 66-593 | 135 ± 60.4 |
| UC-SEC-UC | 1.63E+09 | 74-533 | 134 ± 54.3 |

In pbMEC conditioning medium, the isolated particles were positive for both the EVs markers TSG101 and CD9, while in MAC-T conditioning medium only for TSG101. All lysates were negative for calnexin, ruling out any intracellular contamination [47] (figure 4e).

We isolated detectable amounts of RNA from the pbMECs EVs. Specifically, we obtained on average 11.1 ± 3.5 ng from UF-SEC-UC, 18.8 ± 9.7 ng from UC-SEC-UC (figure 5a). The length of the isolated RNA molecules is displayed in figure 5b and supplementary figure 3, we observed that the amount of 18S and 28S rRNA was higher in UC-SEC-UC. Finally, the isolated EVs contained the miRNA miR-let-7a-5p, that was detected at lower Cts from UC-SEC-UC than from UF-SEC-UC (figure 5c).

**Figure 5.**
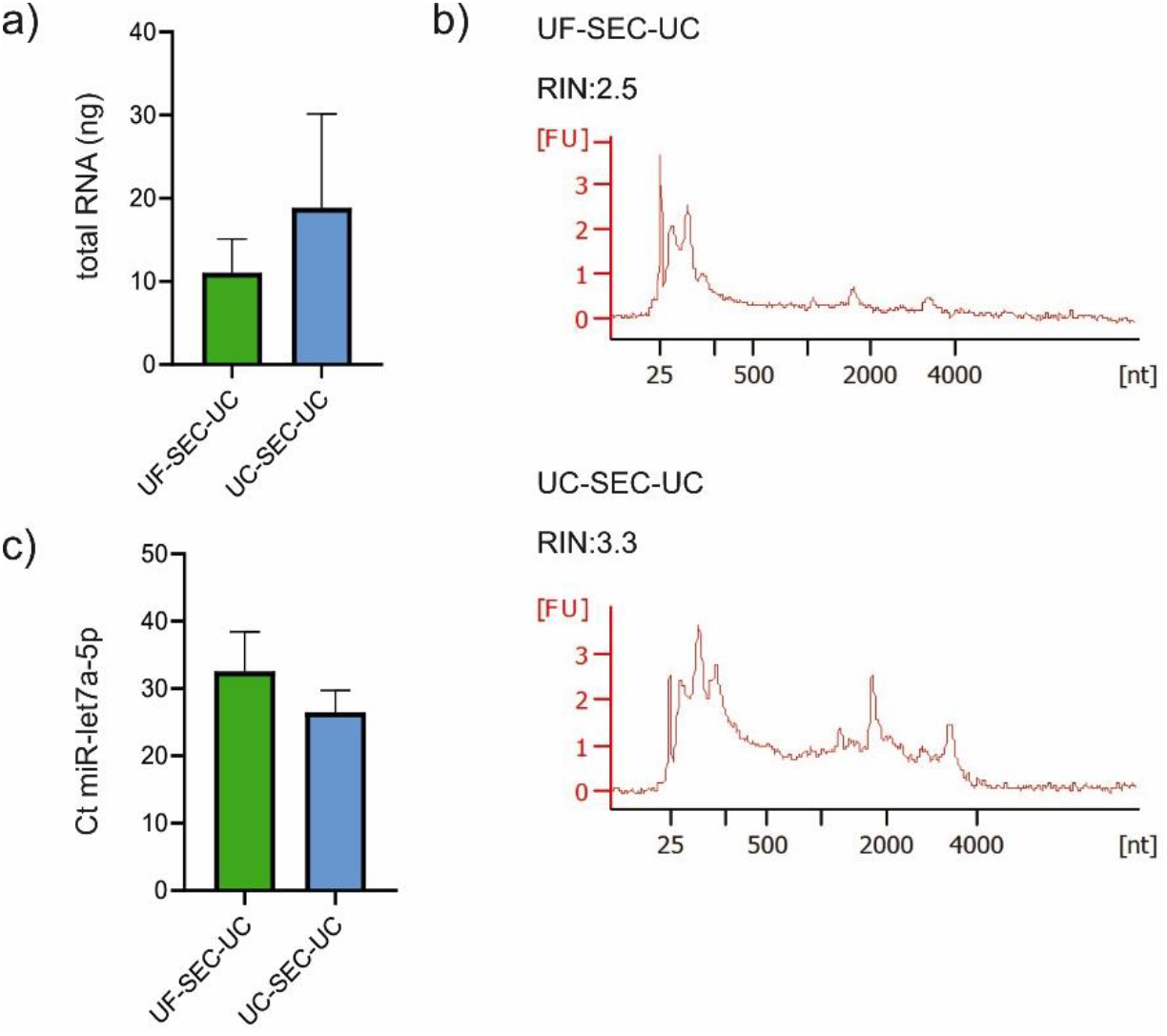
Isolation and characterisation of RNA from pbMEC EV. a) Total amount of RNA from pbMECs EVs isolated with UF-SEC-UC or UC-SEC-UC; b) Representative electropherograms from UF-SEC-UC (top), and UC-SEC-UC (bottom); c) Ct of miR-let-7a-5p from pbMECs EVs isolated with UF-SEC-UC or UC-SEC-UC; a, c) bars show mean and SD of 4 independent experiments.

## DISCUSSION

In the current study, we established for the first time the culture of primary bovine mammary epithelial cells (pbMECs) and MAC-T cells in FBS-free medium to study EVs from the bovine mammary gland in vitro. An important point is that the culture medium is chemically defined, therefore it does not change during the culture and always keeps the same formula, avoiding any possible source of variability given by FBS removal [28, 29] or even different batches of FBS.

Our customized FBS-free medium sustained pbMECs cell growth until confluency and beyond passage 3, as well as the growth of MAC-T cell line. The growth rate of MAC-T was lower than in FBS containing medium. This is in line with previous results, where, however, the culture medium had a different formula [43]. In the primary culture, FBS-free medium promoted the expression of epithelial markers keratin 14 and 18 and the downregulation of the fibroblastic marker vimentin, thereby enriching the population in epithelial cells. In addition, it did not affect the epithelial identity of MAC-T, as keratin 14 and keratin 18 expression levels did not change. Thus, pbMECs and MAC-T cells cultured on FBS free medium were still relatively close to the primary isolated cells at day 0 and FBS-containing medium, respectively.

We also observed a slight downregulation of the differentiation markers PR, PRLR, CSN1S1 and CSN3 already starting at 50% confluency. This trend to de-differentiate might be owed to the culture on two dimensions (2D) on plastic dishes, which by itself promotes de-differentiation [44, 45]. It has been shown that the growth of pbMECs on tri-dimensional (3D) systems supports differentiation [37, 45, 46]. Thus, further studies should focus on 3D FBS-free culture systems.

The most commonly used techniques to isolate EVs from primary MECs cell cultures or breast cancer lines are dual UC [6, 18, 23, 47, 48] and UF with a cut-off of 100 kDa [17], respectively. By combining these methods, size exclusion chromatography and chemical precipitation, we managed to isolate EVs from relatively low amounts (10 mL) of cellular conditioning medium, while in the literature, when stated, the reported starting material is often between 20 and 250 mL [30-32]. The TEM and TRPS measurements excluded any particle contamination in the FBS-free medium, thereby the EVs we observed and analyzed were secreted by the cultured cells. From the eight routes that we tested, only in one route (SEC-UC) we did not isolate any particle after TEM analysis, likely due to the initial low amount of starting material (500 µL). Dual UC gave a high amount of particles but the TEM imaging showed that the matrix was not clean, possible due to the low cleaning steps before the UC. The introduction of more differential centrifugation steps and a SEC step helped to clean the sample likely from proteins in solution and any membranes or content deriving from cell debris and apoptotic bodies. The miRCURY kit used for CP and CP-SEC-UC instead helped to precipitate vesicles but concomitantly generated many undefined aggregates, as the precipitation itself does not distinguish between the types of macromolecules in solution [33, 49]. We obtained a clean sample from UC-SEC-UF, but the particles observed after TEM were few in the whole grid, likely due to the high final volume of the sample (150 µL), collected from the ultrafiltration tube. Both UF-SEC-UC and UC-SEC-UC revealed a better compromise regarding the yield and purity of vesicles. The TRPS measurements confirmed the size ranges observed at TEM and were similar between routes, and both gave a consistent amount of EVs, on the order of 109 -1010 particles/mL. Western blot analysis confirmed that the isolated vesicles were actual EV, bearing both CD9 and TSG101, in line with the results obtained by Zang et al. [17], and from studies on milk EVs [19]. Importantly, the use of a SEC step to separate secreted proteins from EVs would allow analysing the cell response distinguishing the contribution of EVs and secreted proteins. For such purpose, UC-SEC-UC would be more suitable, as the protein-rich fractions are more concentrated, without the initial UF step. On the other hand, if the experimental setup requires the study of EVs only, UF-SEC-UC would be more recommended, as the sample is initially depleted from proteins smaller than 100k kDa.

We were able to extract enough RNA and detect miR-let-7a-5p after using both protocols. MiR-let-7a-5p is a miRNA expressed in the bovine mammary gland tissue [50] and is one of the most abundant exosomal miRNAs found in both human and bovine milk [51]. We observed a higher amount of RNA and miR-let-7a-5p with UC-SEC-UC, however that protocol gave higher contamination from rRNA 18S and 28S, likely pulled down with the first ultracentrifugation step [52].

In conclusion, we demonstrated that the FBS-free medium culture system is a valid tool to study MECs EVs from both primary cells and the MAC-T cell line. We evaluated and compared different EVs isolation protocols from a relatively low amount of starting cell culture medium (10 mL) and we proposed both UC-SEC-UF and UC-SEC-UC are the preferred methods to isolate the highest amount of pure EVs for further downstream analysis. Our results provide an important reference for further studies that aim at analysing MECs EVs in many contexts such as lactation, infection, response to stressors and metabolic challenges.

## Acknowledgements

We thank S. Handschin from the Scientific Center for Optical and Electron Microscopy (ScopeM) of ETH Zurich for his support with TEM. The authors are active participants of the COST Action CA16119 (In vitro 3D total cell guidance and fitness).

## DECLARATIONS

### Authors’ contributions

GS conceptualized the experiments, performed the in vitro cultures, EVs isolation, analysed the data and wrote the manuscript. SEU supervised and financed the project and revised the manuscript. MDS coordinated and supervised the project and revised the manuscript. All authors read, edited and approved the final manuscript.

### Availability of data and materials

Not applicable.

### Ethical approval and consent to participate

Not applicable.

### Consent for publication

Not applicable.

