## Supplementary Figures for "Assessing extracellular vesicles from bovine mammary gland epithelial cells cultured in FBS-free medium"

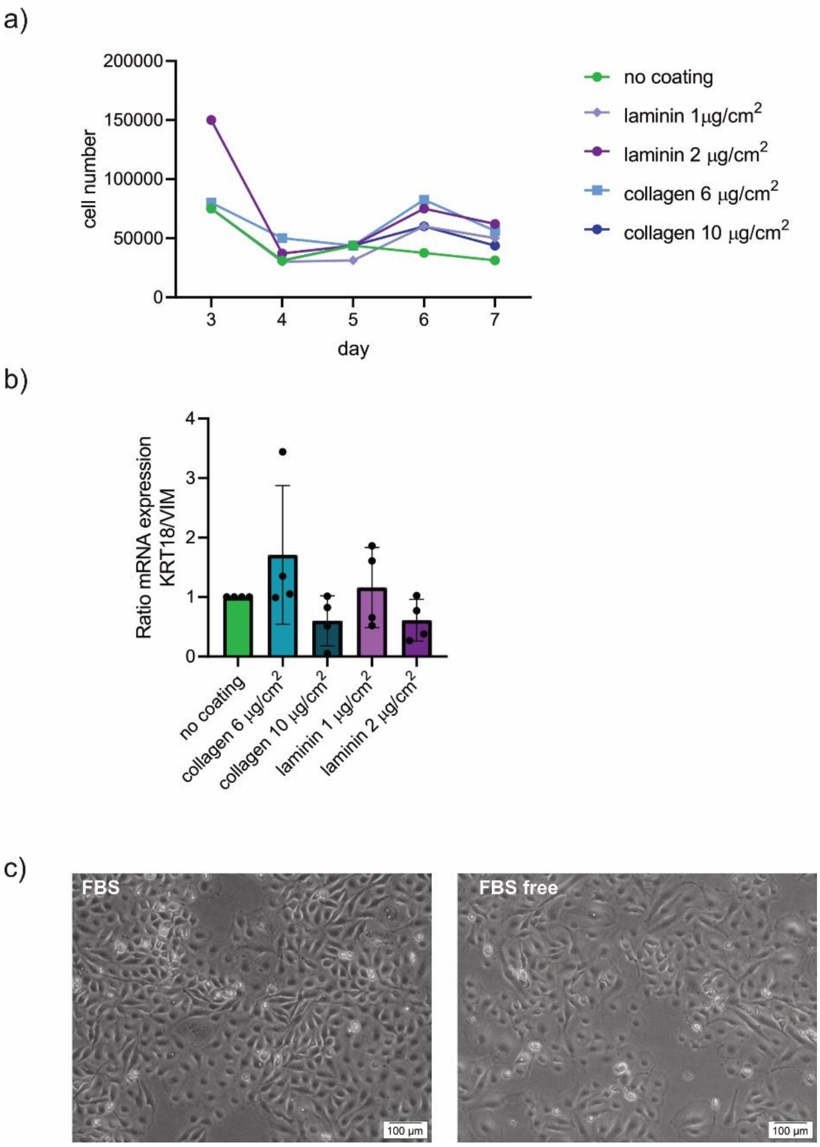


**Supplementary Figure 1.**. a) Growth curve of pbMECs in FBS-free medium. Cells were plated on 6-well multiwell dishes and counted every day from day 3 to day 7; the culture dishes were uncoated or coated with laminin 1 or 2 µg/cm2 or collagen I 6 or 10 µg/cm2; b) Ratio of the mRNA expression of KRT18/VIM of pbMECs at 80% confluence plated in coated or coated wells as in a); c) MAC-T cells grown in FBS 10% or FBS-free medium.

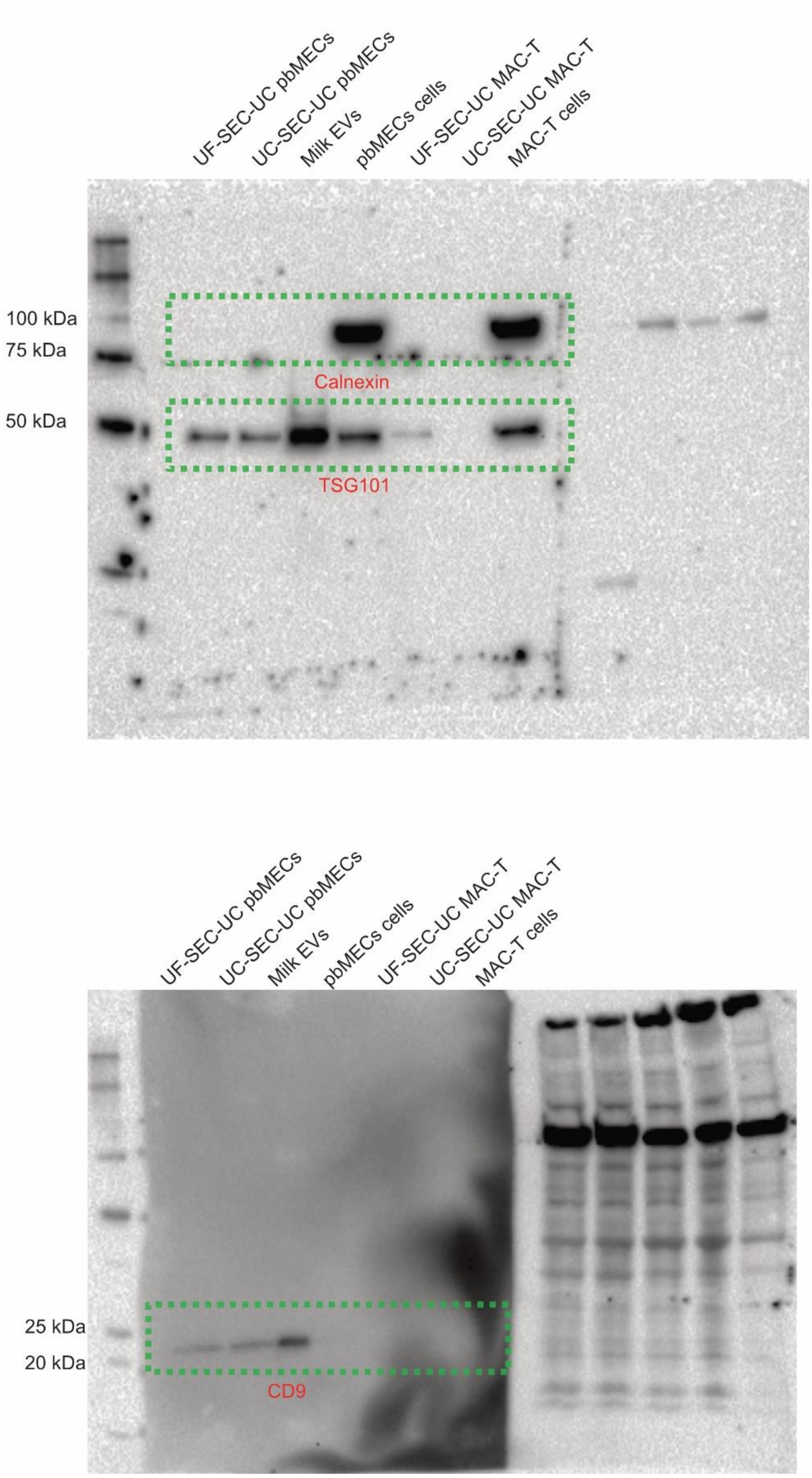


**Supplementary Figure 2.** Full-length western blots shown in figure 4c. The selected areas are in green.

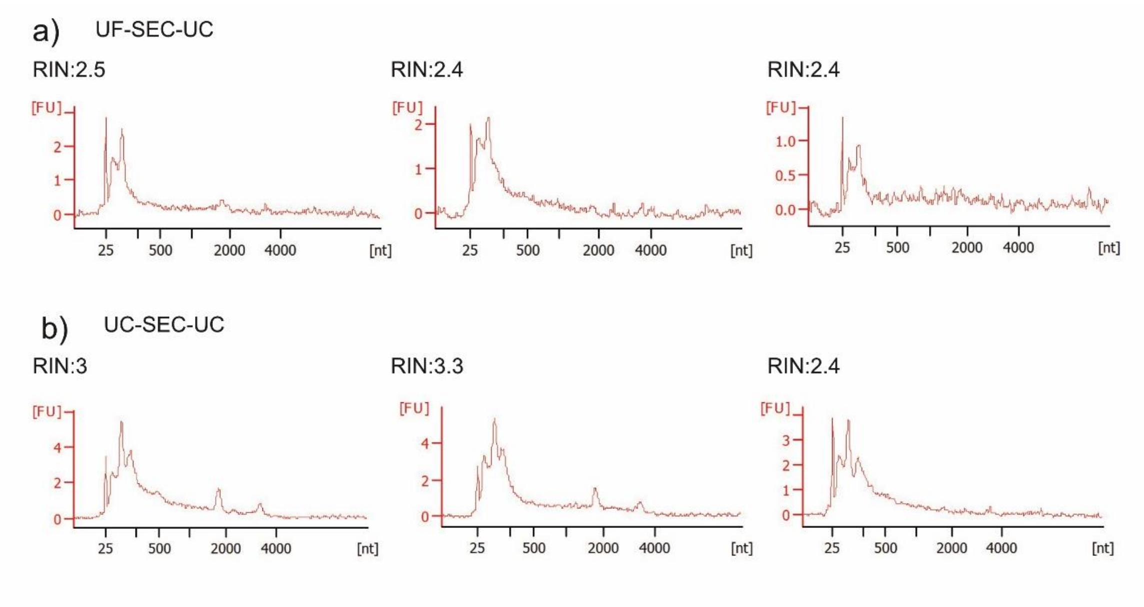


**Supplementary figure 3.** Electropherograms from the Bioanalyzer. a) samples from UF-SEC-UC; b) samples from UC-SEC-UC
